# Ecology of Echinodermata in the Clarion-Clipperton-Fracture Zone (Central Pacific)

**DOI:** 10.1101/2025.06.22.660910

**Authors:** Tanja Stratmann, Erik Simon-Lledó, Marcel T. J. van der Meer, Magdalini Christodoulou, Sven Rossel, Ana Colaço

## Abstract

Abyssal seascapes between 3,000 and 6000 m water depth represent over 50% of the Planet’s surface, but the species, functions, and particularly the life history traits that these ecosystems harbour remain poorly understood. Brittle stars (Ophiuroidea) contribute about one third to the invertebrate megabenthos assemblage between 3,800 m and 4,800 m water depth in the Clarion-Clipperton Fracture Zone (CCZ, Northeast Pacific). Starfishes (Asteroidea) are present in lower densities. In the CCZ, Ophiuroidea are often seen near Xenophyophoroidea and attached to glass sponge (Hexactinellida) stalks. We hypothesize that (1) the observed relationship between Ophiuroidea, Xenophyophoroidea, and Hexactinellida is a predator-prey relationship, where Ophiuroidea feed on foraminifera- and sponge-derived organic matter. (2) Ophiuroidea have a reduced dependency on fresh phytodetritus. (3) Brisingida (order of Asteroidea), often clings to stalks to have easier access to particulate organic matter sinking to the seafloor.

To test these three hypotheses, we combined bulk and compound-specific stable isotope analyses of fauna (Ophiuroidea, Asteroidea) and sediments with the analyses of seafloor images from the eastern CCZ. Faunal specimens and sediments were collected during three research expeditions between 2019 and 2022, and previously collected seabed images were re-analysed to quantify the major behaviours in which Ophiuroidea and Asteroidea engage.

All investigated Echinodermata species had a high trophic level. Phospholipid-derived fatty acids (PLFAs) used as biomarkers suggest that *Silax daleus* consumes sedimentary detritus that is processed by its gut microbiome. *Ophiacantha cosmica* is likely a top consumer or scavenger, *Ophiosphalma glabrum* is an opportunistic omnivore ingesting phytodetritus, bacteria, Crustacea, and Foraminifera, while *Ophiuroglypha* cf. *polyacantha* is a more selective omnivore. *Freyella benthophila* sits mostly on stalks of Hexactinellida and uses this elevated position to catch phytodetritus and zooplankton. *Freyastera* cf. *tuberculata*, in comparison, sits mostly on polymetallic nodules from where it preys upon Crustacea moving on the sediment surface.

This study confirmed the hypothesis that Ophiuroidea in the CCZ are less dependent on phytodetritus than Holothuroidea in the Peru Basin. It was confirmed that Ophiuroidea consume foraminifera- and sponge-derived organic matter, but Brisingida cling to stalks of Hexactinellida to prey upon Crustacea living in the benthic boundary layer.

## 1. Introduction

Abyssal seascapes lay between 3,000 and 6,000 m water depth (Smith et al., 2008) representing the largest ecosystem on Earth (306,595,900 km^2^ area (Harris et al., 2014)) covering 54% of the planet’s surface (Gage and Tyler, 1991)) and 85% of the seafloor (Harris et al., 2014). The best studied abyssal seascapes in the Pacific is the Clarion-Clipperton Fracture Zone (CCZ) between 0°N, 160°W and 23.5°N, 115°W (International Seabed Authority, 2011), which is bound by the Clarion Fracture zone in the north and the Clipperton Fracture Zone in the south. The water depth in the CCZ ranges from about 4,300 to 4,500 m in the shallowest areas of the southeast and central east CCZ, to about 5,400 m in the deepest areas of the northwest CCZ close to Hawaii (Washburn et al., 2021). The bottom water is characterized by temperatures of 1.3 to 2°C, salinities between 34.68 and 34.70, and dissolved oxygen (O_2_) concentrations of 3.6 ml O_2_ l^-1^ in the southeast CCZ to 4.1 ml O_2_ l^-1^ in the northwest CCZ (Washburn et al., 2021). The sediment consists of siliciclastic clay in the north and central CCZ and biogenic calcareous ooze in the south CCZ (Dutkiewicz et al., 2015). Fresh org. C reaches the seafloor as particulate organic carbon (POC) with rates of 1 g C m^-2^ yr^-1^ in the northwest CCZ and 1.74 g C m^-2^ yr^-1^ in the southwest (Washburn et al., 2021). POC flux in the east CCZ can reach up to 2 g C m^-2^ yr^-1^ (Washburn et al., 2021). In large parts of the CCZ, polymetallic nodules, rock concretions of mostly manganese oxides and iron oxy-hydroxides (Hein and Koschinsky, 2014), lay at the sediment surface with densities of 1.5 to 3 kg m^-2^ in the southern CCZ and ∼7.5 kg m^-2^ in the central east CCZ (Washburn et al., 2021), where they provide hard substrate for epi- and infauna (Goineau and Gooday, 2019; Gooday et al., 2020, 2015; Kamenskaya et al., 2013; Lim et al., 2024, 2017; Pape et al., 2021; Simon-Lledó et al., 2019b; Stratmann et al., 2021; Vanreusel et al., 2016).

The invertebrate megabenthos community (i.e., organisms visible on still images, size >1 cm) in the CCZ comprises of hundreds of morphotypes belonging to, at least, 13 different phyla (Simon-Lledó et al., 2023a). Their densities increase with decreasing water depth from (mean ± 95% confidence interval, *CI*) 0.12 (*CI*: 0.06 – 0.14) ind m^-2^ below 4,800 m to 0.87 (*CI*: 0.31 – 1.42) ind m^-2^ above 4,300 m, while diversity (exponential Shannon index, *e^H’^*) decreases with water depth, from 32.7 (*CI*: 22.7 – 42.3) effective taxa below 4,800 m to 21.7 (*CI*: 13.6 – 30.4) in areas above 4,300 m depth (Simon-Lledó et al., 2023b). In the northeast CCZ, three distinct invertebrate megabenthos assemblages were detected that differ in their dominating taxa: Between 3,800 and 4,300 m water depth, the upper-abyssal assemblage is comprised mostly of Alcyonacea (30% relative abundance), Ophiuroidea (22%), Bryozoa (11%), and Demospongiae (7%), whereas the lower-abyssal assemblage between 4,800 to 5,300 m depth consists predominantly of Actiniaria (32%), Holothuroidea (13%), Hexactinellida (13%), and Antipatharia (6%) (Simon-Lledó et al., 2023b). The transitional assemblage at intermediate water depth (4,300 to 4,800 m depth) includes mostly Ophiuroidea (32%), Actinaria (19%), Alcyonacea (9%), and Hexactinellida (6%) (Simon-Lledó et al., 2023b). Overall, while Porifera and Arthropoda megafauna species depict wider distributions across these depth ecotones, Echinodermata and Cnidaria exhibit more restricted dispersal ranges across the CCZ (Simon-Lledó et al., 2025).

Ophiuroidea are very abundant in the CCZ with densities of ∼2,000 ind. ha^-1^ at upper-abyssal depth, ∼1,500 ind. ha^-1^ at intermediate depth, and <40 ind. ha^-1^ at >4,800 m depth (Simon-Lledó et al., 2023b). The species *Ophiosphalma* cf. *glabrum* (Lütken & Mortensen, 1899) is even among the ten most abundant taxa of each depth-dependent CCZ invertebrate megabenthos assemblage (Simon-Lledó et al., 2023b). Besides *O. glabrum*, at least 41 other Ophiuroidea species live in the eastern CCZ (Christodoulou et al., 2020; Eichsteller et al., 2023). These species belong to phylogenetic lineages that fall into three categories based on branch length, with the oldest category including three multi-species clades that are over 70 Mio yr old (Christodoulou et al., 2019).

Asteroidea, in comparison, are less abundant in the CCZ with densities of 411 ind. ha^-1^ at upper-abyssal depth (Uhlenkott et al., 2022), 36 ind. ha^-1^ at intermediate depth (De Smet et al., 2021), and 1 ind. ha^-1^ at lower-abyssal depth (Durden et al., 2021). Hence, they contribute only between 2.8% (upper-abyssal depth), 3.2% (intermediate depth), and 1.7% (lower-abyssal depth) to the CCZ invertebrate megabenthos density, even though the Asteroidea family Paxillosida fam. indet. is among the ten most abundant taxa of the transitional assemblage (Simon-Lledó et al., 2023b). In total, 23 different Asteroidea morphotypes have been reported from the CCZ (Bribiesca-Contreras et al., 2022; Simon-Lledó et al., 2023b). Thus, the Asteroidea richness is considerably lower than the Ophiuroidea biodiversity.

Deep-sea Ophiuroidea often form close commensal relationships with habitat-providing invertebrate megabenthos: For example, in the Atacama Region (Northern Chile, Southeast Pacific) Ophiuroidea cling to the stalk of the Hexactinellida *Caulophacus (Caulophacus) chilensis* Reiswig & Araya, 2014 at 1,300 to, 1,800 m water depth, and they are associated with the Octocorallia *Narella* sp. between 1,800 and 2,000 m depth (de Castro Manso et al., 2018). Further commensal associations between Ophiuroidea and Octocorallia are known from the tropical East Pacific (Exclusive Economic Zone of Columbia), where *Callogorgia* cf. *galapagensis* Cairns, 2018 hosts the Ophiuroidea *Astrodia* cf. *excavata* (Lütken & Mortensen, 1899) between 530 and 670 m water depth (Mejía-Quintero et al., 2021). *Astrodia excavata* may actually spend its whole life cycle with its Octocorallia host (Mejía-Quintero et al., 2021) and research from the western North Atlantic indicates that in some cases, Ophiuroidea age together with their Octocorallia hosts (Mosher and Watling, 2009). When they are very small, the Ophiuroidea *Ophiocreas oedipus* Lyman, 1879 clings to the central stalk of the small Octocorallia *Metallogorgia melanotrichos* (Wright & Studer, 1889) and when the Octocorallia grows older, the Ophiuroidea moves higher up until it sits in the fully developed Octocorallia crown (Mosher and Watling, 2009). In comparison, in the CCZ Ophiuroidea area often observed associated with Xenophyophoroidea (Amon et al., 2016; Christodoulou et al., 2020) and attached to stalks of Hexactinellida (Christodoulou et al., 2020).

Deep-sea Ophiuroidea can have different feeding types or strategies, such as microphagous grazers, scavengers, predators, suspension feeders, or (unselective) omnivores/ opportunistic generalist. Microphagous grazers ingest mostly phytodetritus and a small fraction of sediment (e.g., *Ophiuroglypha lymani* (Ljungman, 1871)) (Dahm, 1999). Scavengers ingest carcasses of e.g., Crustacea, together with a small fraction of phytodetritus and/ or sediment (e.g., *O. lymani*) (Dahm, 1999). Active predators prey upon various prey types and sizes, like Porifera, Ophiuroidea, Bivalvia, Polychaeta, small Crustacea, and/ or Foraminifera (e.g., *Ophiosparte gigas* Koehler, 1922), *Ophiacantha bidentata* (Bruzelius, 1805) (Dearborn et al., 1996; Pearson and Gage, 1984), whereas suspension feeders catch zooplankton and resuspended (particulate) organic matter (P)OM out of the water column (e.g., *Ophiactis abyssicola* (M. Sars, 1861)) (Pearson and Gage, 1984). Omnivores may feed selectively, but most omnivores seem to feed unselectively as opportunistic generalist on a variety of OM, depending on its availability (e.g., *Ophiuroglypha irrorata irrorata* (Lyman, 1878), *Ophiomusa lymani* (Wyville Thomson, 1873), *Ophionotus victoriae* Bell, 1902) (Fratt and Dearborn, 1984; Pearson and Gage, 1984).

Deep-sea Asteroidea show different behaviour depending on whether they belong to the order Brisingida or to other orders, such as Paxillosida or Valvatida. At abyssal depth, Paxillosida may leave impressions in the seafloor (Durden et al., 2020; Miguez-Salas et al., 2024), while transiting quickly between feeding locations with average seafloor tracking rates of 81 to 110 cm^2^ h^-1^ (Durden et al., 2020). During feeding, this Asteroidea order may actively move in the sediment and prey upon sediment infauna (i.e., predatory/ scavenging feeding type) or it may push into the sediment to scope sediment or buried faecal material out into its mouth (i.e., sediment ingester) (Howell et al., 2003; Miguez-Salas et al., 2024). Valvatida may be spongivorous like *Tylaster willei* Danielssen & Koren, 1881 predating upon Demospongiae Tetractinellida gen. indet. between 585 and 722 m at the Langseth Ridge (Arctic Ocean) (Stratmann et al., 2022).

In comparison, deep-sea Brisingida, which are mostly suspension feeders (Howell et al., 2003), use rock outcrops or patches of Scleractinia *Madrepora oculata* Linnaeus, 1758 or *Desmophyllum pertusum* (Linnaeus, 1758) as pedestals in deep-water canyons of the Cantabrian Sea (Northeast Atlantic) (Manjón-Cabeza et al., 2021). Brisingida can also live as epibiota on Octocorallia and in the Caribbean deep sea at 408 m depth, the Brisingida *Novodinia antillensis* (A.H. Clark, 1934) was observed attached to the Octocorallia *Paramuricea* sp. for 14 years (Etnoyer et al., 2022). Hence, similar to Ophiuroidea, Brisingida can form long-lasting mutual relationships. In the CCZ, Brisingida may sit on the sediment or are attached to polymetallic nodules stretching their arms into the water, like *Freyastera* sp., *Freyastera* cf. *tuberculata* (Sladen, 1889), and *Freyella* cf. *benthophila* Sladen, 1889 (Amon et al., 2017; Bribiesca-Contreras et al., 2022). They also cling to the stalk of dead or alive Hexactinellida (Amon et al., 2017; Stratmann et al., 2021).

In this study, we combined (compound-specific) stable isotope analyses of several Ophiuroidea and Asteroidea species with still image analyses of the CCZ seafloor to address the following hypotheses: (1) Ophiuroidea are more abundant in the CCZ than in the Peru Basin (Southeast Pacific), because they are less dependent on fresh phytodetritus. (2) Ophiuroidea are often associated with Xenophyophoroidea and Hexactinellida because they feed on foraminifera- and sponge-derived OM. (3) Brisingida cling to stalks to have an easier access to POM derived from surface waters.

## 2. Materials and methods

### 2.1 Sample collection

During two research cruises to the German and Belgian exploration contract areas in the CCZ in 2019 (expedition SO268 aboard R/V *Sonne* (Haeckel and Linke, 2021)) and 2021 (expedition Mangan2021 aboard M/V *Island Pride* (Vink et al., 2022)), Echinodermata specimens (*n*_total_ = 34; Fig. S1) of the families Freyellidae (Asteroidea), Amphiuridae, Ophiopyrgidae, Ophiosphalmidae, and Ophiacanthidae (all Ophiuroidea) were collected opportunistically with the manipulators of remotely operated vehicles (ROV) Kiel 6000 (Geomar, Germany) and ROV HD14 (Ocean Infinity, Texas, USA) (Table 1; Fig. 1). Further specimens were collected when retrieving a benthic chamber lander and a box corer. Due to opportunistic sampling and the associated logistic constraints, the collection of several species was unbalanced and limited to *n* = 1 or *n* = 2.

**Figure 1.**
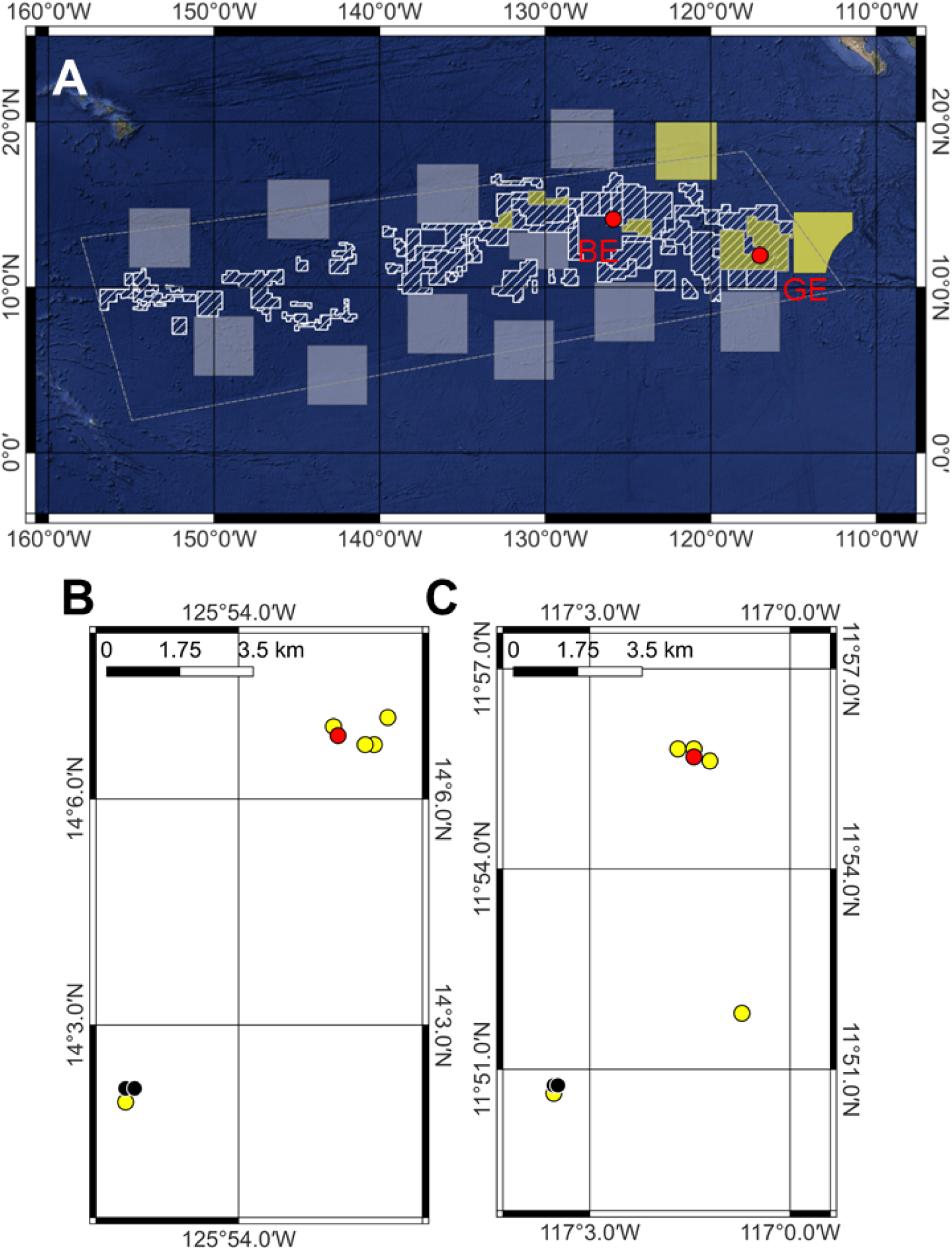
(A) Map of the exploration contract areas (white dashed areas) and areas of particular environmental interest (APEIs; white squares) within the Clarion-Clipperton Fracture Zone (CCZ; white dashed polygon based on the working definition of the CCZ by (Glover et al., 2015)). The transparent yellow areas show APEIs 3 and 12 and exploration license contract areas where seabed imagery had been collected. The two red dots in panel A indicate the Belgian (GSR) and German (BGR) exploration contract areas. (B) Sampling sites in the Belgian exploration contract area. (C) Sampling sites in the German exploration contract area. Color code for panels B and C: Red = sampling site of Asteroidea, black = sediment sampling site, yellow = sampling site of Ophiuroidea. Copyright for shapefiles for the exploration contract areas and APEIs within the CCZ area: ©International Seabed Authority [2007-2020].

**Table 1.** Detailed information about sampling location, date, and species name of all collected Echinodermata.

| Station | Date | Depth (m) | Latitude (°N) | Longitude (°E) | Exploration contract area | Species [Family, Order, Class] |
| --- | --- | --- | --- | --- | --- | --- |
| SO268/1_029_ROV04 | 06/03/2019 | 4090.0 | 11.927 | □117.024 | Germany | *Freyella benthophila* Sladen, 1889 [Freyellidae, Brisingida, Asteroidea] (*n* = 1) |
| SO268/1_034-1_LANDER-03 | 08/03/2019 | 4128.8 | 11.844 | -117.059 | Germany | *Ophiuroglypha* cf. *polyacantha* [Ophiopyrgidae, Ophiurida, Ophiuroidea] (*n* = 1) |
| SO268/1_040_ROV07 | 09/03/2019 | 4134 | 11.845 | -117.059 | Germany | *Ophiosphalma glabrum* (Lütken & Mortensen, 1899) [Ophiosphalmidae, Ophiurida, Ophiuroidea] (*n* = 4), *O.* cf. *polyacantha* (*n* = 1) |
| SO268/1_064_ROV09 | 17/03/2019 | 4506 | 14.112 | -125.870 | Belgium | *Freyastera* cf. *tuberculata* (Sladen, 1889) [Freyellidae, Brisingida, Asteroidea] (*n* = 1), *O. glabrum* (*n* = 6) |
| SO268/1_069_ROV10 | 18/03/2019 | 4504 | 14.112 | -125.871 | Belgium | *O. glabrum* (*n* = 3) |
| SO268/1_077_ROV12 | 20/03/2019 | 4512 | 14.118 | -125.867 | Belgium | *O. glabrum* (*n* = 1), *O.* cf. *polyacantha* (*n* = 1) |
| SO268/2_099_ROV14 | 05/04/2019 | 4127 | 11.863 | -118.863 | Germany | *O. glabrum* (*n* = 2) |
| SO268/2_137_ROV20 | 18/04/2019 | 4496 | 14.111 | -125.872 | Belgium | *O. glabrum* (*n* = 1) |
| SO268/2_154_ROV23 | 23/04/2019 | 4544 | 14.033 | -125.923 | Belgium | *O. glabrum* (*n* = 3) |
| SO268/2_163_ROV25 | 28/04/2019 | 4090 | 11.928 | -117.024 | Germany | *Ophiacantha cosmica* Lyman, 1878 [Ophiacanthidae, Ophiacanthida, Ophiuroidea] (*n* = 1), *O. glabrum* (*n* = 3) |
| IP21_042BC | 27/04/2021 | 4490.0 | 14.116 | -125.879 | Belgium | *O. glabrum* (*n* = 2) |
| IP21_056ROV1 | 04/05/2021 | 4107 | 11.927 | -117.020 | Germany | *O.* cf. *polyacantha* (*n* = 1) |
| IP21_059BC | 05/05/2021 | 4100 | 11.930 | -117.028 | Germany | *Silax daleus* (Lyman, 1879) |
| | | | | | | [Amphiuridae, Amphilepidida,<br>Ophiuroidea] ($n = 1$) |
| IP21_047ROV1 | 02/05/2021 | 4107 | 11.927 | -117.020 | Germany | *O. glabrum* ($n = 1$) |

Aboard the vessels, the asteroid specimens were photographed, subsamples were taken for DNA barcoding, and the specimens were frozen at -20°C. Ophiuroid specimens collected in 2019 were photographed, the arms were separated from the disk for studies of reproduction (Laming et al., 2021), subsample were taken for DNA barcoding, and the remaining arms were frozen at -20. In contrast, in 2021, ophiuroid specimens were photographed, subsamples were taken for DNA barcoding, and the otherwise intact specimens were frozen at -20°C.

Sediment samples (Fig. 1) were taken with a multicorer during research cruise SO295 aboard R/V *Sonne* in 2022 (Haeckel et al., 2023). After retrieval of the multicorer on board, individual cores (*n* = 5) were sliced in three sediment intervals (0 – 2 cm, 2 – 5 cm, 5 – 10 cm sediment layers). Each sediment interval was subsampled for post-cruise analyses of phytopigments, grain size, OM, and phospholipid-derived fatty acids (PLFAs), and stored frozen at -20°C (grain size, OM, PLFAs) and -80°C (phytopigments), respectively.

### 2.3 Still image survey

The behaviour of upper-abyssal depth Echinodermata was recorded from observations captured in seabed imagery collected across multiple sites in the eastern CCZ (APEI 6, APEI 12, UK exploration contract areas, German exploration contract’s eastern subarea, and Tonga exploration contract’s subareas B, C, and D), as compiled in the Abyssal Pacific Seafloor Megafauna Atlas and Database (Simon-Lledó et al., 2023a). Using BIIGLE 2.0 (Langenkämper et al., 2017), behaviours encountered (i.e., Echinodermata on the sediment (Fig. 2A), buried in the sediment (Fig. 2B), under a Xenophyophoroidea (Fig. 2C), under a polymetallic nodule (Fig. 2D), on Scleralcyonacea (class Octocorallia, phylum Cnidaria)/ Hexactinellida stalks (Fig. 2E), under a Hexactinellida (Fig. 2F), or on a polymetallic nodule (Fig. 2G)) were counted for *F. benthophila* sp inc. (AST_053, 14 specimens), *F. tuberculata* sp inc. (AST_002, 73 specimens), *O. glabrum* sp inc. (OPH_010, 4699 specimens), and for another 5174 Ophiuroidea non-identifiable down to morphotype level.

**Figure 2.**
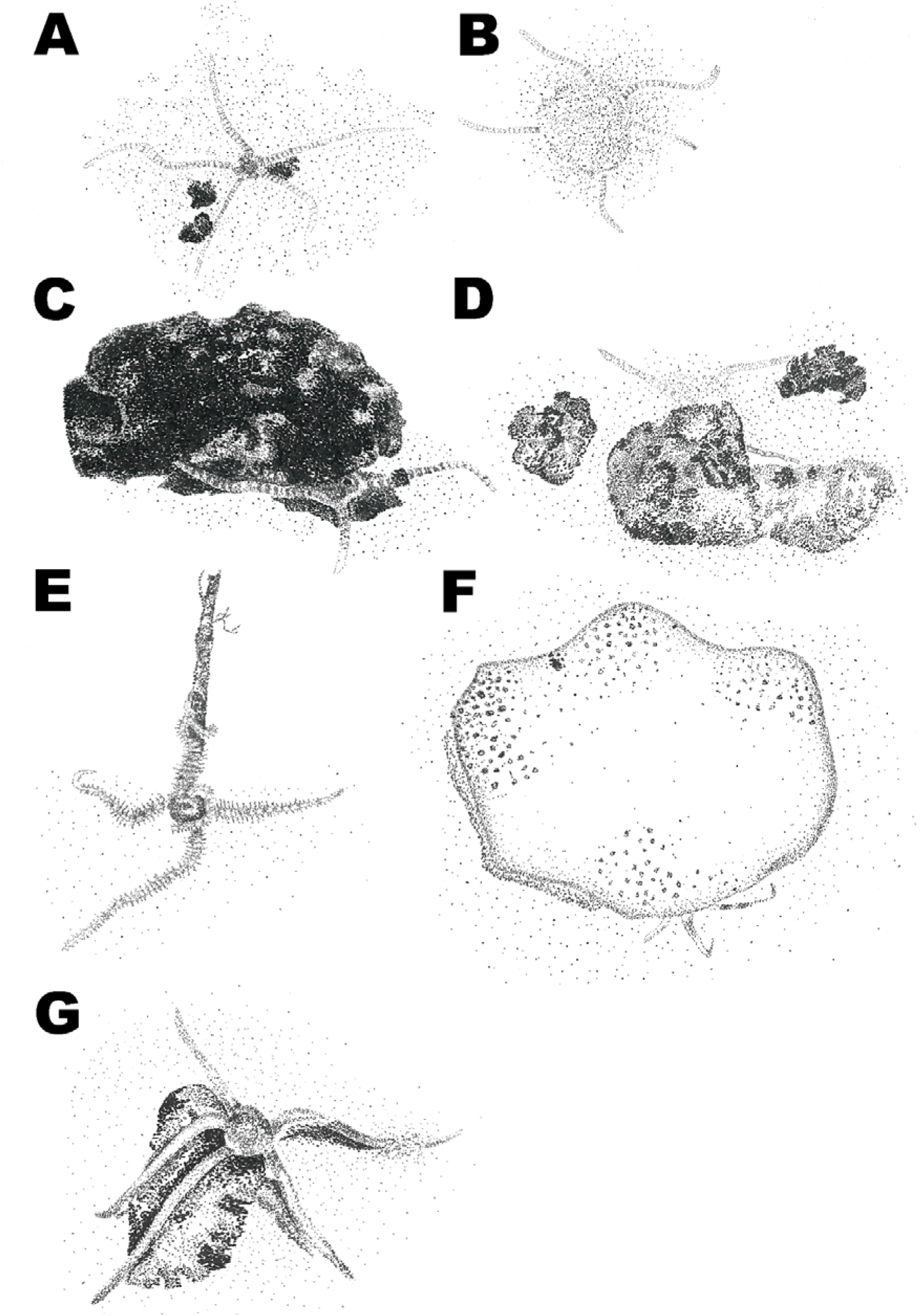
Drawings of the different behaviour types the Echinodermata in the eastern CCZ were engaged in: (A) Ophiuroidea on the sediment, (B) Ophiuroidea buried in the sediment, (C) Ophiuroidea under a Xenophyophoroidea, (D) Ophiuroidea under a polymetallic nodule, (E) Ophiuroidea clinging to a stalk, (F) Ophiuroidea under a Hexactinellida, (G) Ophiuroidea on a polymetallic nodule. Illustration prepared by Tanja Stratmann.

### 2.4 Processing of faunal and sediment samples

#### 2.4.1 Species identification

At Senckenberg am Meer (Wilhelmshaven, Germany), each Ophiuroidea and Asteroidea specimen was initially identified using reverse taxonomy (barcoding), followed by morphological verification. For this purpose, genomic DNA was extracted from arm tissue. DNA extractions were carried out using 30 μl chelex (InstaGene Matrix, Bio-Rad) for 50 min at 56°C, followed by a step at 96°C for 10 min (Estoup et al., 1996). About 2 µl of the resulting DNA-extract was used as template DNA for PCR which was conducted using 12 µl AccuStart (2× PCR master mix, Quantabio), 7.5/9.5 μl molecular grade water, and 0.25/0.5 μl of each primer (20/10 pmol μl^-1^) in a total volume of 20/25 μl. The Ophiuroidea-specific forward primer LCOech1aF1 (5′-TTTTTTCTACTAAACACAAGGATATTGG-3′) (Layton et al., 2016) with the universal reverse primer II: HCO2198 (5′-TAAACTTCAGGGTGACCAAAAAATCA-3′) (Folmer et al., 1994) were used for the amplification. Amplification was carried out in a thermocycler with the following settings: Initial step at 94°C for 3 min, denaturation step at 94°C for 30 s, annealing at 43°C for 75 s, elongation at 72°C for 60 s, and a final elongation at 72°C for 3 min. Denaturation, annealing and elongation were carried out in 40 cycles. The success of all reactions was checked on a 1%-agarose gel; all PCR-products that produced a band were sent to Macrogen Europe (Amsterdam, Netherlands), for sequencing on an Applied Biosystems 3730XL sequencer. A negative control was used in all PCR runs. Resulting sequencing reads were assembled in *Geneious R9* (version 9.1.7) and checked for contamination (e.g., bacteria, fungi) using the basic local alignment search tool (BLAST) (Altschul et al., 1997). A list with all genbank accession codes for the identified specimens is presented in Table S1.

#### 2.4.2 Bulk analyses

In the laboratories of NIOZ-Texel (Den Hoorn, Texel, Netherlands) and of the University of the Azores (Horta, Portugal), Echinodermata and sediment samples were freeze-dried and ground to fine powder with mortar and pestle. Total C (TC), total N (TN), δ^13^TC, and δ^15^TN contents of Echinodermata and sediments were measured in not acidified samples with a Thermo Flash EA 1112 elemental analyzer (EA; Thermo Fisher Scientific, USA) coupled to a DELTA V Advantage Isotope Ratio Mass Spectrometer (IRMS; Thermo Fisher Scientific) at NIOZ-EDS (Yerseke, Netherlands). Organic C and δ^13^org. C content of Echinodermata and sediments were measured with the same EA-IRMS after sample acidification with 20 µl 2% HCl. δ^13^C and δ^15^N data were normalized against the isotope reference materials USGS40 (Qi et al., 2003) and USGS41a (Qi et al., 2016) as described in (Sharp, 2017). The stable isotope data are presented in δ notation relative to Vienna Pee Dee Belemnite (VPDB; δ^13^C) and air (δ^15^N), respectively.

Particle sizes of freeze-dried, sieved (<1 mm) sediment was measured by laser diffraction in a Malvern Mastersizer 2000 (Malvern Instruments, UK).

Phytopigments were extracted from ∼1.5 g dry mass (DM) freeze-dried sediment with 10 ml 90% acetone + β-Apo-8′-carotenal (APO) as described in (Rios-Yunes et al., 2023). Pigment concentrations (in µg g^-1^ DM sediment) were measured via a high-performance liquid chromatography (HPLC) with a LC-04 (Shimadzu Co., Japan) using a Waters C8 column (length: 150 mm, inner diameter: 4.6 mm, particle size: 3.5 µm, pore size: 0.01 µm; Waters Corporation, USA) connected to a Symmetry C8 Sentry Guard Cartridge (length: 20 mm, inner diameter: 3.9 mm, particle size: 5 μm, pore size: 100 Å; Waters Corporation) following the approach by (Zapata et al., 2000).

#### 2.4.3 Fatty acid analyses

At NIOZ-Texel, PLFAs from freeze-dried Echinodermata powder (∼0.03 – 0.07 g DM) and freeze-dried ground sediment (1.9 – 3.2 g DM) were extracted following a modified version of the Bligh and Dyer extraction (Bligh and Dyer, 1959; Boschker, 2008) as described in detail in (Stratmann et al., 2023), though chloroform had been replaced by dichloromethane (DCM). In brief, total lipids were extracted with the Bligh & Dyer extraction and fractionated into different lipid classes over a silicic-acid column. The PLFAs dissolved in the DCM fraction were derivatized into fatty acid methyl esters (FAMEs) via mild alkaline transmethylation and FAMEs were separated on a BPX70 column (length: 50 m length, inner diameter: 0.32 mm, film thickness: 0.25 μm; SGE Analytical Science, Australia) as described in detail in (Stratmann et al., 2024). C concentration of individual FAMEs (in µmol C-PLFA g^-1^ DM sediment) and their δ^13^C_VPDB_ values (in ‰) were measured on a Delta V Advantage IRMS (Thermo Scientific, USA) coupled to a Trace 1310 GC (Thermo Scientific) with a Isolink II interface coupled to a Conflo IV (Thermo Scientific). Only those fatty acids were considered, that were present in at least 25% of the sediment samples or 15% of the Echinodermata specimens.

A list with fatty acids that can serve as biomarkers is presented in Table 2.

**Table 2.** Fatty acids used as biomarkers of potential food sources of Echinodermata from the Clarion-Clipperton Fracture Zone. Abbreviations: ARA = arachidonic acid (C20:4ω6), DHA = docosahexaenoic acid (C22:6ω3), EPA = eicosapentaenoic acid (C20:5ω3).

| Fatty acid | Main source of fatty acid/ feeding type | Reference |
| --- | --- | --- |
| **Bacteria** |  |  |
| *i*-C14:0, *i*-C15:0, *ai*-C15:0, *i*-C16:0, and C18:1ω7 *cis* | Marine bacteria | (Middelburg et al., 2000) |
| C14:0, C16:0, C16:1ω7 *cis*, 16:1ω5, C16:1ω8 *cis*, and 18:1ω7 *cis* | Chemosynthetic bacteria from intertidal sediments/ cold seeps | (Boschker et al., 2014; Van Gaeve et al., 2009b) |
| C14:0, C16:0, C16:1ω7 *cis/trans*, and 18:1ω7 *cis* | Sulfur and ammonium oxidizing bacteria | (Blumer et al., 1969; Knief et al., 2003; Lipski et al., 2001; Sakata et al., 2008; Van Gaeve et al., 2009a; Zhang et al., 2005) |
| *i*-C17:1ω7, *ai*-C17:1ω7, C17:1ω8 *cis*, cy-C17:0/C17:1ω6 *cis*; C16:1ω5; 10Me-C16:0, cy-C18:0 *i*-C17:1ω7 *cis*; *i*-C15:1ω7 *cis*; *i*-C19:1ω7 *cis* | Sulfate reducing bacteria | (Boon et al., 1977; Elvert et al., 2003; Oude Elferink et al., 1998; Taylor and Parkes, 1983) |
| C16:1ω5 *trans*, C16:1ω6 *cis*, C16:1ω8 *cis*, C16:1ω7, C18:1ω8 *cis*, C18:1ω11 *cis* | Methanotrophic bacteria | (Bowman et al., 1993; Nichols et al., 1987) |
| C16:1ω7, C18:1ω7 | Chemosynthetic, sulfide oxidizing bacteria | (Colaço et al., 2007) |
| C18:1ω13 | Methylotrophic bacteria | (Colaço et al., 2007) |
| (a)i-PLFA | Gram-positive bacteria | (Zelles, 1997) |
| cy-PLFA | Gram-negative bacteria | (Zelles, 1997) |
| Primary producers |  |  |
| C16:4ω1, C16:1ω7, and EPA;<br>$\frac{C16:1\omega7}{C16:0}$-ratio>1 or $\frac{DHA}{EPA}$-ratio<1 | Diatoms | (Budge and Parrish, 1998;<br>Dalsgaard et al., 2003;<br>Kharlamenko et al., 1995;<br>Parrish et al., 2005) |
| C16:4ω1, C16:1ω7, and EPA;<br>$\frac{C16:1\omega7}{C16:0}$-ratio<1 or $\frac{DHA}{EPA}$-ratio>1 | Dinoflagellates | (Budge and Parrish, 1998;<br>Dalsgaard et al., 2003;<br>Parrish et al., 2005) |
| Consumers |  |  |
| C18:1ω9, C20:1ω9, C20:1ω11,<br>C22:1ω11 | Copepods, Amphipods | (Auel et al., 2002;<br>Dalsgaard et al., 2003; Falk-Petersen et al., 1987;<br>Kattner et al., 2012) |
| PLFAs with $\geq 24$ Cs | Sponges | (de Kluijver et al., 2021) |
| ARA; 22:5ω5; $\frac{EPA}{ARA}$-ratio | Agglutinated<br>foraminifera | (Kharlamenko, 2018; Larkin<br>et al., 2014) |
| Feeding types |  |  |
| $\frac{C18:1\omega9}{C18:1\omega7}$-ratio>1 | Carnivory index | (Graeve et al., 1997) |

### 2.5 Data analyses

Bacterial biomass (*BB*; µg C g^-1^ DM sediment) was calculated following (Middelburg et al., 2000) as:

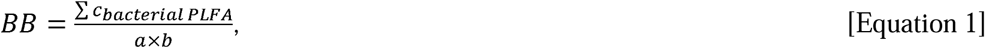

where *c_bacterial PLFA_* is the concentration of the bacteria-specific PLFAs *i*-C14:0, *i*-C15:0, *ai*-C15:0, *i*-C16:0, and C18:1ω7, *a* is the average concentration of PLFAs in bacteria (i.e., 0.056 g C-PLFA g^-1^ C biomass), and *b* is the average fraction of bacteria-specific PLFAs to total PLFAs in sediments (i.e., 0.28) (Middelburg et al., 2000).

Trophic positions (*TP*s) of the different Echinodermata species was estimated with the “oneBaseline” model of the package *tRophicPosition* (“Bayesian trophic position estimation with stable isotopes”; version 0.80) (Quezada-Romegialli et al., 2018) in *R* (version 4.4.3) (R-Core Team, 2025). This model calculates *TP* as follows:

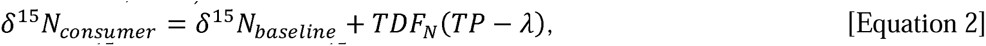

where δ^15^*N_consumer_* is the δ^15^ value of the individual asteroid and ophiuroid species, *δ*^15^*N_baseline_* is the *δ*^15^ value of the sediment averaged across all three layers, *TDF*_N_ is the trophic discrimination factor for N (i.e., 3.4‰ (Post, 2002)), and λ is the trophic position of the sediment (i.e., 1.0).

*TPs* were grouped into trophic levels (*TLs*) following (Yunda-Guarin et al., 2022): TPs >1 and ≤2 are primary consumers (low *TL*), *TPs* >2 and ≤3 are secondary consumers, such as omnivores (intermediate *TL*), and TP >3 are top consumers and scavengers (high *TL*).

Corrected standard ellipse area (*SEA_c_*; ‰^2^) as proxy for isotopic niche width (*INW*) was calculated with the package *SIBER* (“Stable Isotope Bayesian Ellipses in R”; version 2.1.9) (Jackson et al., 2011) in *R*.

All data are presented as mean ± standard error (SE).

## 3. Results

### 3.1 Biogeochemical composition of sediment

Sediments (*n* = 5) in the eastern CCZ contained of 0.58 ± 0.04 % TC, 0.54 ± 0.05% org. C, and 0.11 ± 0.00% TN in the upper 2 cm of sediment, 0.51 ± 0.03% TC, 0.45 ± 0.04% org. C, and 0.10 ± 0.00% TN in the 2 to 5 cm layer, and 0.46 ± 0.03% TC, 0.38 ± 0.03% org. C, and 0.09 ± 0.00% TN in the 5 to 10 cm layer. The C/N ratio decreased from 5.59 ± 0.35 in the 0 to 2 cm layer to 4.75 ± 0.27 in the 5 to 10 cm layer. The sediment had δ^13^org C values of - 18.5 ± 0.29‰ (0 – 2 cm layer), -18.2 ± 0.08‰ (2 – 5 cm layer), and -18.3 ± 0.13‰ (5 – 10 cm layer) and δ^15^TN values of 12.3 ± 0.11‰ (0 – 2 cm layer), 12.4 ± 0.17‰ (2 – 5 cm layer), and 12.0 ± 0.18‰ (5 – 10 cm layer) (Fig. 3).

**Figure 3.**
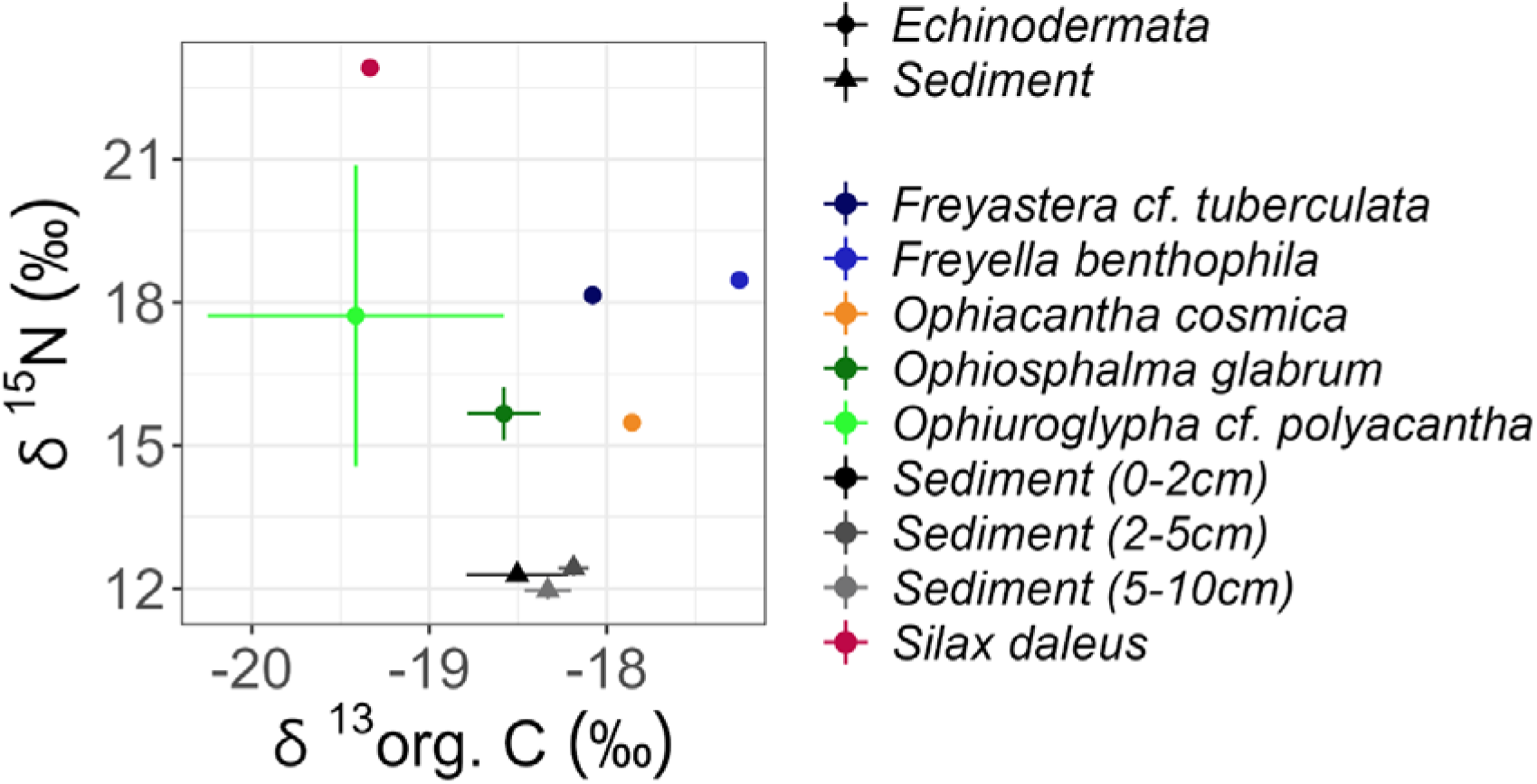
Isotopic composition of organic C (δ^13^org. C, ‰) and N (δ^15^N, ‰) of Echinodermata and sediment from the Clarion-Clipperton Fracture Zone. Error bars indicate 1 SE.

Most of the sediment consisted of silt (<63 µm grain size fraction; 88.1 ± 1.61% in the 0 – 2 cm layer to 89.3 ± 1.08% in the 2 – 5 cm layer), followed by very fine sand (62.5 – 125 μm grain size fraction; 6.48 ± 0.73% in the 0 – 2 cm layer to 6.95 ± 0.89% in the 5 – 10 cm layer) and fine sand (125 – 250 μm grain size fraction; 3.10 ± 0.46% in the 5 – 10 cm layer to 3.67 ± 0.51% in the 0 – 2 cm layer). The median grain size ranged from 7.24 ± 0.69 µm in the 5 – 10 cm layer to 9.11 ± 0.82 µm in the 0 – 2 cm layer.

Phytopigment concentrations were below the detection limit (chlorophyll-*a*: <0.03 µg g^-1^ DM sediment, pheophytin: <0.2 µg g^-1^ DM sediment, alloxanthine: 0.01 µg g^-1^ DM sediment, diatoxanthin: <0.09 µg g^-1^ DM sediment) in any of the sediment layers.

The surface sediment layer (0 – 2 cm) contained 0.48 ± 0.10 µmol C-PLFA g^-1^ DM sediment, whereas the 2 – 5 cm and 5 – 10 cm layers contained 0.36 ± 0.07 µmol C-PLFA g^-1^ DM sediment and 0.22 ± 0.07 µmol C-PLFA g^-1^ DM sediment. The PLFAs C16:0, C18:0, C18:1ω7, and C20:1ω9 *cis* contributed 79.4 ± 5.24% to the total PLFA concentration in the 0 – 2 cm layer, whereas C16:0, C18:0, and C20:4ω6 (arachidonic acid, ARA) contributed 72.7 ± 9.65% and 81.9 ± 9.63%, respectively, to the total PLFA concentrations in the 2 – 5 cm and 5 – 10 cm layers (Fig. S2A).

*BB* was 9.65 ± 4.45 µg g^-1^ DM sediment in the 0 – 2 cm layer, 3.90 ± 2.11 µg g^-1^ DM sediment in the 2 – 5 cm layer, and 1.33 ± 0.96 µg g^-1^ DM sediment in the 5 – 10 cm layer.

### 3.2 Trophic position, trophic level, and isotopic niche width of Echinodermata

Trophic positions of Echinodermata lay in the high trophic level range (Table 3). *Ophiosphalma glabrum* (Lütken & Mortensen, 1899 had the lowest mean *TP* (3.0 ± 0.003), whereas *Silax daleus* (Lyman, 1879) had the highest mean *TP* (4.3 ± 0.04; Table 3).

**Table 3.** Trophic position (*TP*) and level (*TL*) of all Echinodermata species investigated in this study. Data are presented as mean ± SE.

| Species [Class] | *TP* | *TL* |
| --- | --- | --- |
| *Freyella benthophila* [Asteroidea] | $3.9 \pm 0.04$ | High TL |
| *Freyastera* cf. *tuberculata* [Asteroidea] | $4.0 \pm 0.04$ | High TL |
| *Ophiuroglypha* cf. *polyacantha* [Ophiuroidea] | $3.7 \pm 0.02$ | High TL |
| *Ophiosphalma glabrum* [Ophiuroidea] | $3.0 \pm 0.003$ | Intermediate/<br>high TL |
| *Ophiacantha cosmica* [Ophiuroidea] | $3.6 \pm 0.04$ | High TL |
| *Silax daleus* [Ophiuroidea] | $4.3 \pm 0.04$ | High TL |

*SEA_c_* of *O. glabrum* was 9.39‰^2^ (95% *CIs*: 5.84 – 13.2‰^2^) and *SEA_c_* of *O*. cf. *polyacantha* was 37.5‰^2^ (95% *CIs*: 3.14 – 84.0‰^2^), and the two isotopic niches overlapped by 56.2%.

### 3.3 Biochemical composition of Echinodermata

Asteroidea tissue contained 16.2 ± 2.88% TC, 8.41 ± 2.25% org. C, and 2.24 ± 0.54% TN (*n* = 2) and Ophiuroidea tissue had 13.5 ± 1.49% TC, 2.92 ± 0.61% org. C, and 0.90 ± 0.22% TN (*n* = 32). *F. benthophila* (*n* = 1) and *S. daleus* (*n* = 1) had the lowest TC (13.3 and 12.6%), org. C (6.16 and 2.07%), and TN (1.70 and 0.47%) contents in Asteroidea and Ophiuroidea, respectively, whereas *F.* cf. *tuberculata* (*n* = 1) and *O. glabrum* (*n* = 26) had the highest TC (19.0 and 13.6%), org. C (10.7 and 2.91%), and TN (2.79 and 0.91%) contents. Mean δ^13^TC-values of Asteroidea and Echinodermata were -11.0 ± 1.25 and -6.32 ± 0.85‰ [min: -12.3‰, *F.* cf. *tuberculata* (*n* = 1), max: -5.86 ± 0.57‰, *Ophiuroglypha* cf. *polyacantha* (*n* = 4)], mean δ^13^org. C-values were -17.7 ± 0.41 and -18.7 ± 0.80‰ [min: - 19.3‰, *S. daleus* (*n* = 1), max: -17.3‰, *F. benthophila* (*n* = 1)], and mean δ^15^N-values were +18.3 ± 0.16 and +16.1 ± 2.49‰ [min: +15.5‰, *Ophiacantha cosmica* Lyman, 1878 (*n* = 1), max: +22.9‰, *S. daleus (n = 1)*] (Fig. 3).

### 3.4 Phospholipid-derived fatty acid composition of Echinodermata

The PLFA composition (Fig. 4A, Fig. S1B) of Echinodermata differed strongly between species. Between 31.8% (*F.* cf. *tuberculata*) and 69.1% (*F. benthophila*) of the PLFAs detected in Asteroidea and Ophiuroidea consisted of mono-unsaturated fatty acids (MUFA) (Fig. S1). The other PLFA classes found were saturated fatty acids (SFA; 41.1 ± 2.51%), branched-chain fatty acids (BCFA; 12.4 ± 1.67%), highly unsaturated fatty acids (HUFA, i.e., fatty acids with ≥4 double bonds; 5.03 ± 1.17%) and polyunsaturated fatty acids (PUFA, i.e., fatty acids with ≥2 double bonds; 4.29 ± 0.77%). Compared to the average PLFA composition across all Echinodermata investigated in this study, *F. benthophila* (6.74%) and *S. daleus* (25.9%) contained less SFA. Tissue of *F.* cf. *tuberculata* (2.71%) and *O.* cf. *polyacantha* (4.45%) consisted of less BCFA than the average Echinodermata, whereas *F. benthophila* tissue (16.8%) included more BCFA compared to the average Echinodermata. MUFA were present in higher percentages in *F. benthophila* (69.1%), *O.* cf. *polyacantha* (45.4%), and *S. daleus* (56.2%) compared to the average Echinodermata in this study, whereas *O. glabrum* (33.1%) and *F.* cf. *tuberculata* (31.8%) had a lower relative contribution.

**Figure 4.**
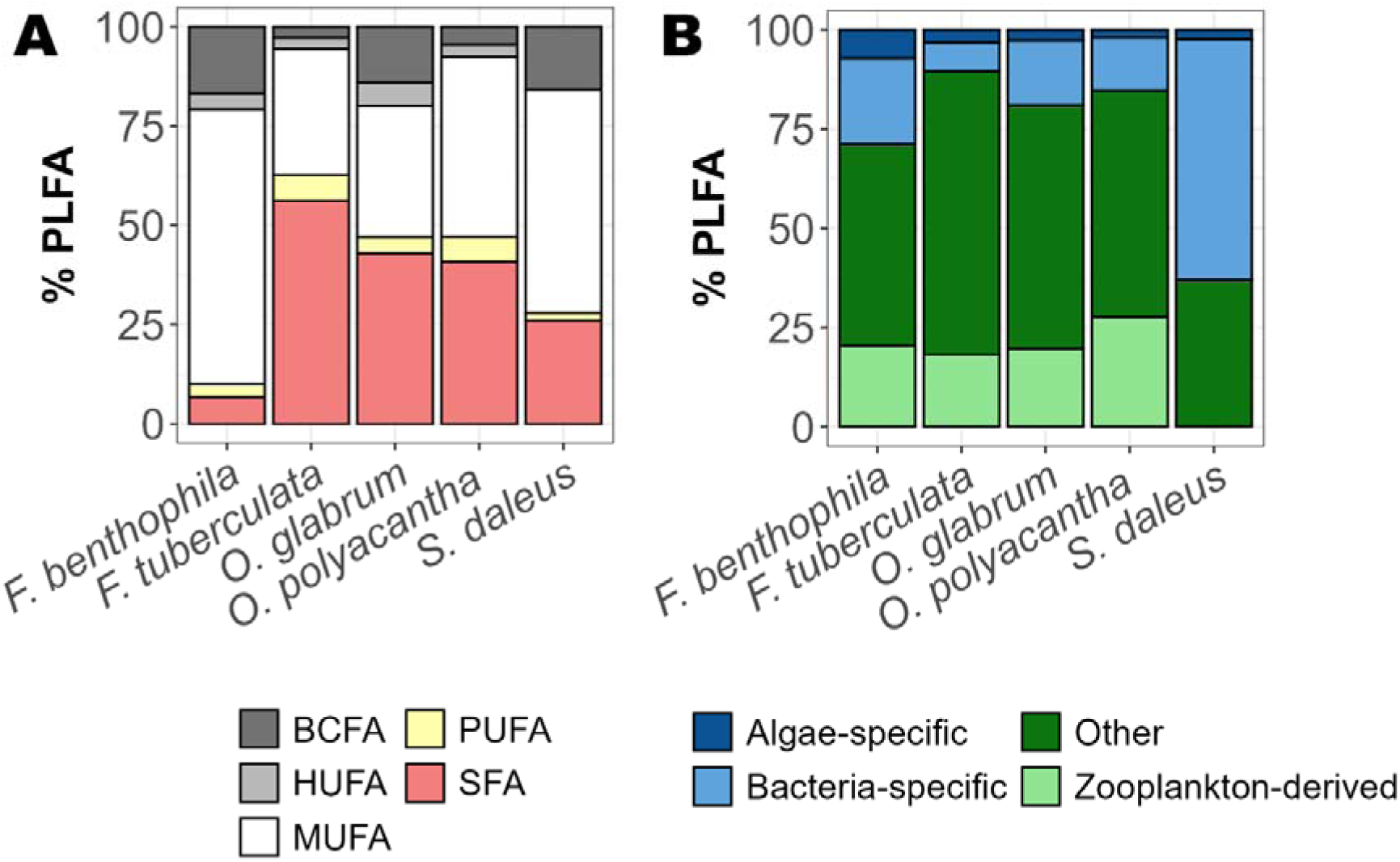
Contribution (%) of (**A**) individual phospholipid-derived fatty acid (PLFA) classes to the total concentration in Echinodermata and (**B**) of individual PLFA categories to total PLFA concentrations. Abbreviations of PLFA classes: BCFA, branched-chain fatty acids; HUFA, highly unsaturated fatty acids; MUFA, mono-unsaturated fatty acids; PUFA, polyunsaturated fatty acids; SFA, saturated fatty acid.

Most PLFAs in the different Echinodermata were unspecific (range: 37.1 – 71.3%), followed by bacteria-specific PLFAs (range: 7.15 – 60.4%). Up to 27.6% of the PLFAs were zooplankton-derived and between 1.97% and 7.22% of the PLFAs originated from primary producers, like algae (Fig. 4B).

The carnivory index was 0.00 for *S. daleus* (*n* = 1), 0.74 for *F. benthophila* (*n* = 1), 2.50 for *F.* cf. *tuberculata* (*n* = 1), 1.53 ± 0.54 for *O.* cf. *polyacantha* (*n* = 3), and 3.96 ± 1.05 for *O. glabrum* (*n* = 16).

Stable C isotope values (δ^13^C) of PLFAs across all investigated Echinodermata species ranged from -29.9‰ (5^th^ percentile) to -11.6‰ (95^th^ percentile) (mean ± SE: -23.6 ± 0.21‰; median: -23.5‰). When separated by species, *F. benthophila* had the most depleted PLFAs (mean ± SE: -25.8 ± 2.46‰; median: -23.1‰; 5^th^ – 95^th^ percentiles: -36.8‰ – -19.7‰ ) and *S. daleus* had the least depleted PLFAs (mean ± SE: -21.0 ± 0.74‰; median: -20.6‰; 5^th^ – 95^th^ percentiles: -25.4‰ – -19.0). The most depleted PLFAs were *i*-C14:0 (-37.1‰, n = *1*) and C18:2ω6,9t (-36.2‰, n = *1*) in *F. benthophila* and C20:3ω3,6,9c (mean ± SE: - 31.0 ± 2.83‰; median: -29.8‰) and *ai*-C15:0 (mean ± SE: -30.4 ± 4.00‰; median: -30.6‰) in *O. glabrum* (Fig. 5).

**Figure 5.**
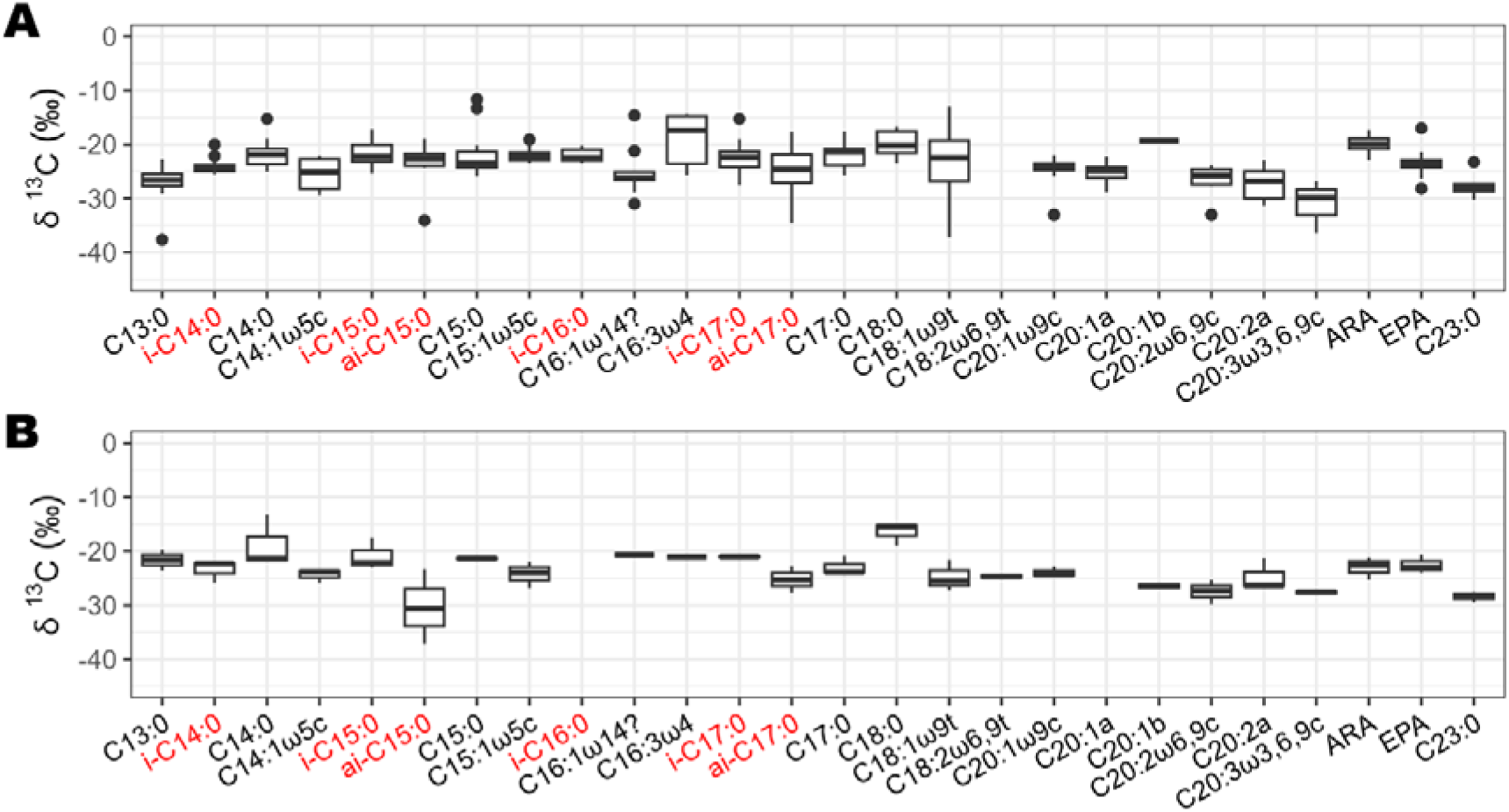
Isotopic composition (δ^13^org. C, ‰) of PLFAs in (**A**) *O. glabrum* (*n* = 19) and (**B**) *O.* cf. *polyacantha* (*n* = 3). PLFA presented in red are bacteria-specific PLFAs.

### 3.4 Behavior of Echinodermata based on still-image surveys

A total of 9960 upper-abyssal depth Echinodermata were behaviourally classified into seven categories based on seabed imagery from the eastern CCZ (Fig. 2, Fig. 6). *Ophiosphalma glabrum* sp inc. (4699 specimens) and unidentified Ophiuroidea (5174 specimens) were markedly more abundant than the Asteroidea *F. benthophila* sp inc. (14 specimens) and *F. tuberculata* sp inc. (73 specimens).

**Figure 6.**
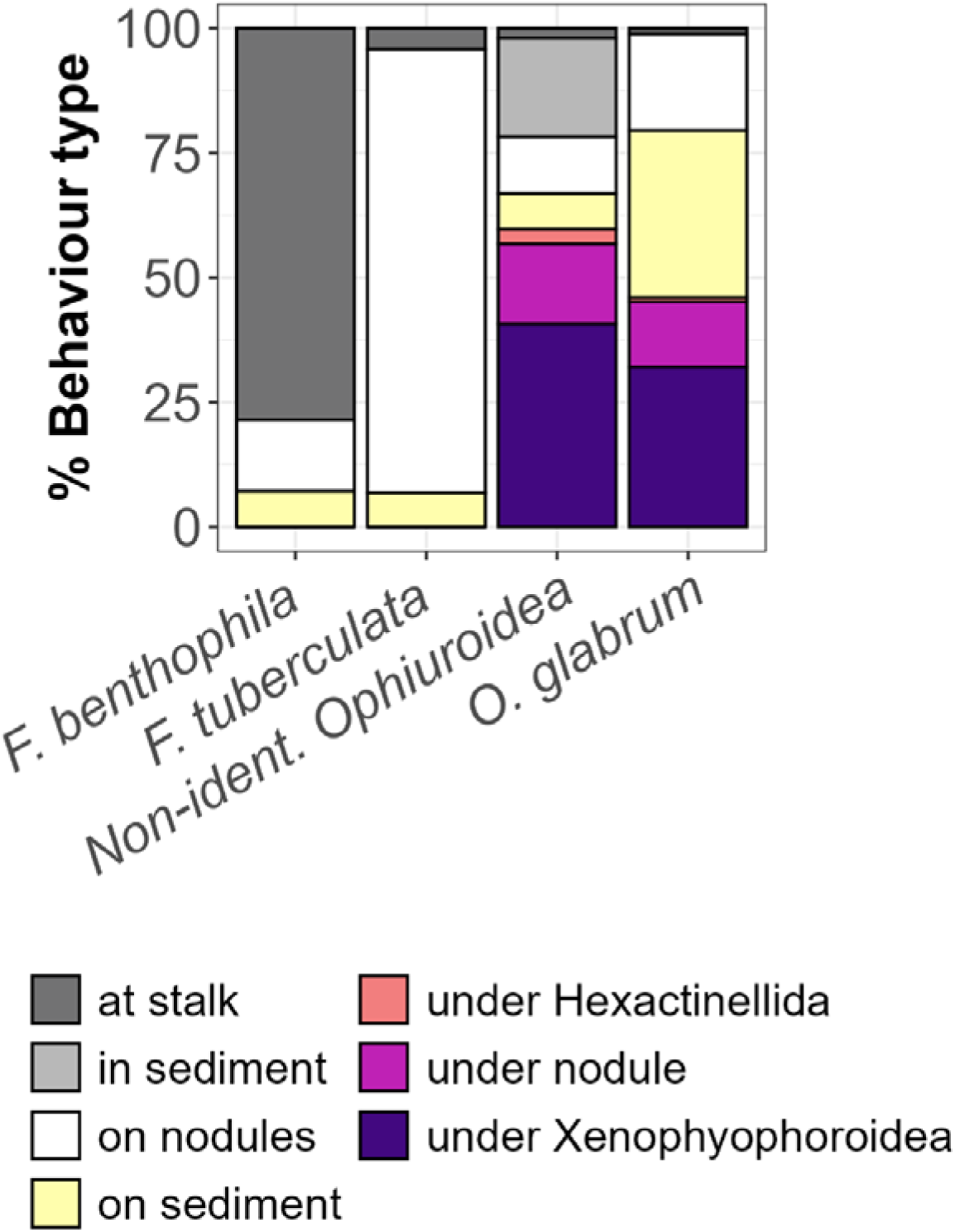
Types of behaviour that Echinodermata showed in the eastern CCZ.

*Freyella benthophila* sp inc. showed a strong preference for occupying elevated positions, with 78.6% observed on stalks of Hexactinellida (Fig. 6); surprisingly, *F. benthophila* sp inc. was never observed at Scleralcyonacea stalks. In comparison, *F. tuberculata* sp inc. was predominantly found on polymetallic nodules (89.0%; Fig. 6). The three *F. tuberculata* sp inc. specimens, that were observed clinging to a stalk, were all attached to *Hyalonema* sp. sp inc. (HEX_002, (Simon-Lledó et al., 2023a)) and lived close-by in the BGR exploration license area.

*Ophiosphalma glabrum* sp inc. exhibited more varied habits: 33.5% were found on sediment, 32.0% under Xenophyophoroidea, and 19.4% on polymetallic nodules (Fig. 6). *O. glabrum* sp inc. observed clinging to stalks were mostly found attached to the base of the main branch of Scleralcyonacea stalks. Individual Ophiuroidea buried in sediment could not be identified to morphotype level owing to limited visibility, likely resulting in an underrepresentation of *O. glabrum* sp inc. in this category. Conversely, Ophiuroidea resting on the sediment surface could often be confidently assigned to OPH_010 (Simon-Lledó et al., 2023a), reducing the pool of unidentified specimens.

Unidentified Ophiuroidea also showed behavioural diversity, with 40.7% under Xenophyophoroidea, 19.7% buried in sediment, and 16.1% under polymetallic nodules (Fig. 6). Notably, a high proportion of unidentified Ophiuroidea at APEI 12 exhibited a burrowing behaviour, with 47.0% of non-identified specimens found with their bodies submerged in sediment, considerably higher than the 4 – 11% range observed across the other locations. Based on arm morphology, many of these specimens are suspected to be *O. glabrum* sp inc. Aside from this site-specific trend, no major spatial differences were observed in the relative proportions of behavioural categories across the region.

Overall, behaviours involving concealment (under polymetallic nodules, Hexactinellida, or Xenophyophoroidea) were common, accounting for 53.2% of all Ophiuroidea records, particularly under giant Foraminifera, the most common behaviour recorded (36.6%).

## 4. Discussion

The class Ophiuroidea is one of the dominant metazoan invertebrate megabenthos classes in the CCZ, but its ecological role is largely understudied. For instance, it is unknown why Ophiuroidea are almost five times more abundant than Holothuroidea at upper-abyssal depths in the CCZ (Simon-Lledó et al., 2023b), whereas both classes have relatively comparable densities in the Peru Basin (Ophiuroidea: 200 – 270 ind. ha^-1^ (Simon-Lledó et al., 2019a); Holothuroidea: 151 – 241 ind. ha^-1^ (Stratmann et al., 2018c)). Reasons could be feeding and diet preferences of Ophiuroidea, and their behaviour.

Here, we decipher feeding strategies and types of several Ophiuroidea and Asteroidea species and describe their behaviour to assess the research hypotheses that (1) Ophiuroidea in the CCZ are less dependent on fresh phytodetritus than Holothuroidea in the Peru Basin, that (2) Ophiuroidea feed foraminifera- and sponge-derived OM, and that (3) the members of the Asteroidea order Brisingida climb up the stalk of Hexactinellida to have easier access to POM from surface waters.

### 4.1 Diets of Ophiuroidea

Information about Ophiuroidea feeding types or strategies may be obtained from gut content analyses (e.g., (Dahm, 1999; Pearson and Gage, 1984), stable isotope analyses (e.g., (Kharlamenko et al., 2013; Yang et al., 2020)), and biomarker (i.e., PLFA) analyses (e.g., (Drazen et al., 2008)). Combined stable isotope and PLFA analyses unravelled the diet preferences and feeding types of four species Ophiuroidea from the CCZ.

#### Silax daleus

(family Amphiuridae) has an estimated trophic position of 4.3 which corresponds to the trophic level of top consumers or scavengers (Yunda-Guarin et al., 2022). However, the PLFA extracted from the tissue of *S. daleus* suggest otherwise: A carnivory index of 0.00 implies this species is not a carnivore (i.e., predator or scavenger), and the dominance of bacteria-specific PLFAs (60.4% of total PLFA) and a 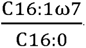-ratio of 0.28 (i.e., <1) indicate that *S. daleus* is a microphagous grazer grazing upon bacteria and dinoflagellate-derived phytodetritus.

The bacteria could be sedimentary bacteria and/ or be part of the gut microbiome of *S. daleus*. Sedimentary bacteria in the upper 5 cm of sediment have a higher biomass in the eastern CCZ (Belgian exploration contract areas: 548 ± 274 mg C m^-2^, German exploration contract areas: 413 ± 314 mg C m^-2^, both this study; UK exploration contract areas: 277 ± 52 mg C m^-2^ – 530 ± 104 mg C m^-2^ (Sweetman et al., 2019)) than in the same sediment layer in the Peru Basin (294 ± 42 mg C m^-2^ (Stratmann et al., 2018b)). Hence, *S. daleus* would have access to more bacteria than Ophiuroidea living in the Peru Basin, but the composition of bacteria-specific PLFAs in *S. daleus* tissue differs from the composition of these PLFAs in the sediment. The CCZ sediment only contained (*a*)*i-*C15:0, *i*-C16:0, and C18:1ω7 as bacteria-specific PLFAs (Table S2), whereas *S. daleus* tissue included additionally the bacteria-specific PLFAs (*a*)*i*-C17:0 (Table S3). As the latter PLFAs are lacking in the sediment, they likely originate from the Ophiuroidea microbiome. Unfortunately, we did not study the gut microbiome of any Ophiuroidea or Asteroidea specimens in this study, but (Dong et al., 2021) showed that shallow-water Ophiuroidea (50 m water depth) from the Yellow Sea (Northwest Pacific) host 56 Bacteria phyla belonging to 596 genera in their gut. Therefore, we assume that *S. daleus* consumes sedimentary detritus which is processed by its gut microbiome, like the Holothuroidea species *Benthodytes* sp. or *Mesothuria* sp. in the Peru Basin (Stratmann et al., 2023).

#### Ophiacantha cosmica

(family Ophiacanthidae) has an estimated trophic position of 3.6. Unfortunately, we lack the PLFA profile of this species to confirm the corresponding trophic level. Therefore, we tried to infer its potential feeding type from the PLFA profiles and gut content of other *Ophiacantha* species (i.e., *Ophiacantha* sp. and *O. bidentata*). At the abyssal ‘Station M’ (Northeast Pacific), the PLFA composition of *Ophiacantha* sp. contains 24.3% PLFAs that are biomarkers for copepods and amphipods (C18:1ω9, C20:1ω9, C20:1ω11, C22:1ω11 (Auel et al., 2002; Dalsgaard et al., 2003; Falk-Petersen et al., 1987; Kattner et al., 2012)) (Drazen et al., 2008). In comparison, at bathyal depth in the Rockall Trough (Northeast Atlantic), *O. bidentata* ingests Foraminifera, Polychaeta, Crustacea, and POM (Pearson and Gage, 1984). Hence, *O. cosmica* in the CCZ is likely indeed a top consumer and/ or scavenger.

#### Ophiosphalma glabrum

(family Ophiosphalmidae) has an estimated trophic position of 3.0 which means it has an intermediate to high trophic level and could be an omnivore or top consumer/ scavenger. A predatory/ scavenging feeding strategy was confirmed by the carnivory index of 3.96 and by the presence of biomarkers for copepods, amphipods (i.e., C18:1ω9 *cis*/*trans*, C20:1ω9 *cis*), and Foraminifera (i.e., 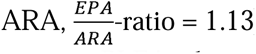) which contribute 23.1% to total PLFAs. *O. glabrum* tissue also contains PLFAs that are biomarkers of primary producers (i.e., C16:1ω7, and EPA) and 16.3% of the PLFAs are bacteria-specific PLFAs. Hence, this Ophiuroidea species is in fact an omnivore ingesting diatom-derived phytodetritus (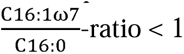), bacteria, and prey or scavenges upon Crustacea and Foraminifera.

#### Ophiuroglypha cf. polyacantha

(family Ophiopyrgidae) has an estimated trophic position of 3.7 and therefore a high trophic level, which indicates *O.* cf. *polyacantha* might be an omnivore or top consumer/ scavenger. Since the biomarkers present in tissue of *O.* cf. *polyacantha* and *O. glabrum* are very similar (Table S3, Fig. 4B, Fig. S2), *O.* cf. *polyacantha* is also an omnivore consuming mostly zooplankton and Foraminifera, and to a lesser degree bacteria and diatom-derived phytodetritus. However, its *INW* is broader (37.5‰^2^) than the niche width of *O. glabrum* (9.39‰^2^).

In the marine environment, a broader *INW* may be an indicator for a dietary specialist, whereas a dietary generalist may have a smaller *INW* (Cummings et al., 2012). Isotopic niche width may also vary with spatial scale, whereupon at finer scale (<1 – 10 km lag) the *INW* is smaller than at larger scale, and the change in *INW* stabilizes at mesoscale (>20 km lag) (Reddin et al., 2018). Thus, *O.* cf. *polyacantha* may be a more specialized omnivore than *O. glabrum* that is likely a more opportunistic/ general omnivore, and/ or *O glabrum* forages at a more confined spatial scale compared to *O.* cf. *polyacantha*.

These results confirmed the hypothesis that Ophiuroidea in the CCZ are not very dependent on phytodetritus: The PLFA composition of all Ophiuroidea species investigated in this study contained less than 10% algae-specific PLFAs, whereas the Holothuroidea species from the Peru Basin that (Stratmann et al., 2023) studied, contained on average 14.4 ± 2.77% (min: 0.09%, max: 41.4%) of these specific PLFAs. In fact, the authors identified two trophic groups of Holothuroidea and both groups had a diet that was more or less rich in phytodetritus. In contrast, the most abundant Ophiuroidea species in the eastern CCZ, *O glabrum*, is an opportunistic omnivore that feeds to a large portion Foraminifera and zooplankton. The latter could be bentho-pelagic zooplankton, but it is also possible that *O glabrum* scavenges (gelatinous) zooplankton carcasses that reach the abyssal seafloor stochastically during mass deposition events, like pyrosome carcasses in the Peru Basin (Hoving et al., 2023) or carcasses of the red crab *Grimothea planipes* (Stimpson, 1860) in the CCZ (Simon[Lledó et al., 2023).

Our diet analyses partly confirmed the hypothesis that Ophiuroidea consume foraminifera- and sponge-derived OM. While the PLFA profiles of *O. glabrum* and *O.* cf. *polyacantha* contained PLFAs typical for Foraminifera, no sponge-specific PLFAs were detected in any investigated Ophiuroidea (or Asteroidea) species. Hence, the 36.6% of Ophiuroidea specimens that resided partly below Xenophyophoroidea (Fig. 2A) prey either actively upon Xenophyophoroidea or they ingested detritus that these produced. Hiding of Ophiuroidea below Hexactinellida (Fig. 2B), however, is not directly linked to spongivory.

### 4.2 Diets of Asteroidea

The order Brisingida has been studied in different oceans which allows us to assess population-specific differences in diets of *F. benthophila*.

#### Freyella benthophila

has an estimated trophic position of 4.0 which corresponds to a high trophic level, like a top consumer/ scavenger or omnivore. In fact, 20.5% of the PLFAs present in *F. benthophila* are biomarkers for zooplankton, whereas 7.22% of the PLFAs are algae-specific PLFAs. Based on the carnivory index of 0.74, this Asteroidea species likely ingests less consumers (e.g., Crustacea, Foraminifera) than the general omnivore *O. glabrum*. It is also possible that *F. benthophila* consumes more degraded, carrion-derived OM, such as detritus derived from Foraminifera, (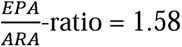) or faecal pellets from zooplankton (Wilson et al., 2013).

At the abyssal Porcupine Seabight (Northeast Atlantic), the suspension feeding *Freyella elegans* (Verrill, 1884) ingested almost exclusively copepods, and the large concentration of biomarkers for primary producers was interpreted to originate from phytodetritus that the copepods had ingested prior to being preyed upon by *F. elegans* (Howell et al., 2003). *F. benthophila* in the eastern CCZ could also target copepods, that had fed mostly diatom-derived phytodetritus (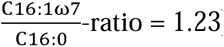), and pelagic Foraminifera while suspension feeding.

#### Freyastera cf. tuberculata

has an estimated trophic position of 4.0 indicating that this species may be a top consumer/ predator or omnivore. The high trophic level is confirmed by a carnivory index of 2.50 and the presence of the zooplankton biomarkers C18:1ω9 *cis* and C20:1ω9 *cis* that contribute 18.2% to total PLFAs. No Foraminifera biomarkers were detected in *F.* cf. *tuberculata* suggesting that this species captures selectively Crustacea when suspension feeding.

### 4.3 Behaviour of Echinodermata

The behaviour of *F. benthophila* (clinging to Hexactinellida stalks) is in line with its feeding type, i.e., suspension feeder targeting (bentho-)pelagic zooplankton and phytodetritus. In fact, it is interesting that *F. benthophila* seems to only climbs up Porifera stalks and not Scleralcyonacea stalks. Deep-sea members of the class Octocorallia are carnivores and prey upon zooplankton (Rodkina and Dautova, 2025), so not clinging to this taxonomic group, but to filter-feeding Hexactinellida indicates that *F. benthophila* may avoid direct competition for food.

*F.* cf. *tuberculata* specimens were mostly sitting on top of polymetallic nodules, which, together with the results of the PLFA analyses, suggests, that this species preys less upon (bentho-) pelagic zooplankton and more upon Crustacea that ‘walk over the seafloor’ instead of swim.

Overall, we could not confirm the hypothesis that Brisingida cling to Porifera stalks for easier access to POM. Instead, our results from diet and image analyses suggested that Brisingida adapted this behaviour because they prey upon copepods or other Crustacea living in the benthic boundary layer.

The large behavioural diversity of *O. glabrum* matches its omnivorous feeding type. The most interesting observation, however, is the dominance of burial behaviour of unidentified Ophiuroidea (likely mostly *O. glabrum* specimens) at APEI 12, compared to the behaviour of (unidentified) Ophiuroidea living in other parts of the eastern CCZ. APEI 12 is the shallowest area investigated in this study (mean water depth: 3940 m (Simon-Lledó et al., 2023b)) and lays above the carbonate compensation depth (CCD; ∼4400 to ∼4800 m water depth (Berger et al., 1976)). The CCD influences the species distribution and biogeography of benthic Foraminifera (Gooday et al., 2021; Gooday and Jorissen, 2012), as calcareous Foraminifera have difficulties to maintain their tests at water depth below the CCD where the tests can dissolve (Corliss and Honjo, 1981). Below the CCZ, the Foraminifera assemblage consists mostly of agglutinated Foraminifera and Foraminifera with organic walls (Saidova, 1965) in (Gooday and Jorissen, 2012) (Shi et al., 2020). Though no data are available for Foraminifera species composition at APEI 12, it is very likely that calcareous Foraminifera are part of the sedimentary infauna at this site.

Here, we propose that *O. glabrum*, which feeds on Foraminifera in the Belgian and German exploration contract areas, buries in the sediment at APEI 12 to prey upon calcareous Foraminifera, which are likely absent in the deeper areas of the eastern CCZ. As a result, a lower number of *O. glabrum*/ non-identified Ophiuroidea specimens bury at the other investigated sites where the Foraminifera community may be different. This stresses the opportunistic feeding of *O. glabrum* that seems to ingest whatever is available and switches diets depending on where a specific specimen lives.

## 5. Conclusion

Our study suggests that omnivorous Ophiuroidea may be more abundant at shallow-abyssal depth in the CCZ than deposit-feeding Holothuroidea, because they may outcompete Holothuroidea at feeding. The CCZ is more oligotrophic than the Peru Basin (Jahnke, 1996) and receives less fresh phytodetritus (CCZ: chlorophyll-*a* and phaeopigment concentrations <detection limit, this study; Peru Basin: chlorophyll-*a* concentration: 0.09 μg ml^-1^ sediment, phaeopigment concentration: 0.23 μg ml^-1^ sediment (Vonnahme et al., 2020)). Hence, omnivores, like Ophiuroidea, that can opportunistically switch their diets and ingest less phytodetritus when this is sparsely available, have an advantage over more specialized deposit feeders, like Holothuroidea, that more selectively take up phytodetritus (Hudson et al., 2003; Stratmann et al., 2023, 2018b; Wigham et al., 2003). In the Peru Basin, in comparison, where the density of Ophiuroidea and Holothuroidea is comparable, sufficient phytodetritus reaches the seafloor to support a relatively large biomass of Holothuroidea (de Jonge et al., 2020; Stratmann et al., 2018c, 2018a), and omnivorous Ophiuroidea can reduce competition for food by adjusting diet preferences based on food availability.

## Acknowledgements

We thank captain, crew, and chief scientists of the research expeditions SO268 (Dr. Peter Linke, Geomar, Germany) aboard R/V Sonne and Mangan2021 aboard M/V Island Pride (Dr. Annemiek Vink, BGR, Germany) for their excellent support at sea. We also acknowledge the invaluable help of the teams of ROV HD14 (Ocean Infinity, Austun, USA) and ROV Kiel 6000 (Geomar, Kiel, Germany). Jurian Brasser, Peter van Breugel (both NIOZ-EDS), Jort Ossebar, and Ronald van Bommel (both NIOZ-MMB) are thanked for technical assistance during sample processing.

Research leading to these results has received funding from JPI Oceans—Ecological Aspects of Deep Sea Mining project under NWO-ALW grant 856.18.003. TS was further supported by the Dutch Research Council NWO (NWO-Rubicon grant no. 019.182EN.012, NWO-Talent program Veni grant no. VI.Veni.212.211, NWO Open Competition Domain Science – XS grant no. OCENW.XS24.2.193). ESL received funding from the Ramón y Cajal grant RyC2023-043275-I, funded by the MCIN/AEI/10.13039/501100011033 and the European Union Next Generation EU/PRTR. SV was funded through the 2020-2021 Biodiversa and Water JPI joint call for research projects, under the BiodivRestore ERA-NET Cofund (grant no. 101003777), with the EU and German Federal Ministry of Research (BMBF) (03LW0173). AC’s work received national funds through the FCT – Foundation for Science and Technology, I.P., under the project UIDB/05634/2025 and UIDP/05634/2025, and in the scope of the CEEC contract CEECIND/00101/2021.

This is publication number XY of Senckenberg am Meer Metabarcoding and Molecular Laboratory.

## Author contributions

**T Stratmann** conceived the ideas and designed the methodology; **T Stratmann** and **A Colaço** collected Echinodermata specimens, and **E Simon-Lledó** collected and annotated the still images. **M Christodoulou** and **S Rossel** performed the molecular analyses for species identification, **T Stratmann** performed all biochemical analyses. **T Stratmann** and **MTJ van der Meer** processed all PLFA data. **T Stratmann** led the writing of the manuscript, all authors contributed critically to the drafts and gave their final approval for publication.

## Statement on inclusion

The study was conducted in areas beyond national jurisdiction (‘the Area’) of the central Pacific. Therefore, no local partners were contacted and involved in this study, as no clear local partners exist.

## Data availability statement

All data presented in this study are available at Pangaea (XX), on Genbank (XX), and are deposited in the barcode of life data system (BOLD) in the project ‘XX’.

